# Conditional ERK3 overexpression cooperates with PTEN deletion to promote lung adenocarcinoma formation in mice

**DOI:** 10.1101/2021.07.11.451860

**Authors:** Sreeram Vallabhaneni, Jian Liu, Marion Morel, Francesco J. Demayo, Weiwen Long

**Author notes:** These authors contributed equally to this work (Sreeram Vallabhaneni, Jian Liu).

## Abstract

Extracellular signal-regulated kinase 3 (ERK3), also known as MAPK6, belongs to the atypical mitogen-activated protein kinases (MAPKs) subfamily. In comparison with the well-studied classical MAPKs ERK1 and ERK2, much less is known about the cellular and molecular actions of ERK3. Accumulating studies have revealed the upregulation of ERK3 expression and suggested an important role for ERK3 in promoting tumor cell growth and invasion in multiple cancers, in particular lung cancer. However, it is unknown whether or not ERK3 plays a role in spontaneous tumorigenesis. To determine the role of ERK3 in lung tumorigenesis, we created a conditional ERK3 transgenic mouse line in which ERK3 transgene expression is controlled by Cre recombinase. By crossing with a lung tissue-specific CCSP-iCre mouse line, we have found that conditional ERK3 overexpression cooperates with PTEN deletion to induce the formation of lung adenocarcinomas (LUADs). Mechanistically, ERK3 overexpression stimulates activating phosphorylations of ERBB3 and ERBB2 by upregulating SP1-mediated gene transcription of NRG1, a potent ligand for ERBB3/ERBB2. To our knowledge, our study is the first revealing a bona fide tumor-promoting role for ERK3 using genetically engineered mouse models. Together with previous findings showing important roles of ERK3 in cultured cells and in xenograft lung tumor model, our findings corroborate that ERK3 acts as an oncoprotein in promoting LUAD development and progression.

## Introduction

Extracellular signal-regulated kinase 3 (ERK3) is a member of the atypical mitogen-activated protein kinases (MAPKs) [1]. It is considered an atypical MAPK in that ERK3 signaling is not organized as classical three-tiered kinase cascades, and its kinase domain harbors a Ser-Glu-Gly (SEG) activation motif instead of the Thr-Xaa-Tyr (TXY) activation motif shared by the classical MAPKs such as ERK1 and ERK2 [1–2]. While still much less is known about the molecular actions of ERK3 signaling in cancers in comparison with the well-studied ERK1/2 signaling, recent years have seen a considerable gain of our understanding of the roles of ERK3 in cancer development. On the one hand, ERK3 has been shown to promote cancer cell growth and migration in culture conditions and tumor growth and metastasis in xenograft mouse models of different human cancers, including lung cancer [3–6], head and neck cancer [7], and breast cancer [8–10]. On the other hand, the inhibitory roles for ERK3 in tumor cell growth and/or migration have also been reported in several other types of cancers, including melanoma [11], non-melanoma skin cancer [12], hepatocarcinoma [13], and intrahepatic cholangiocarcinoma [14]. Taken together, these studies suggest that ERK3 plays either tumor-promoting or tumor-suppressive roles depending on specific cancer type. Several different molecular mechanisms underlying the cancer-promoting role of ERK3 have been proposed, such as activating SRC-3/PEA3-mediated MMP gene transcription in lung cancers and breast cancer [3] and c-Jun/AP1-mediated IL-8 expression in colon cancer cells and MDA-MB231 breast cancer cells [9]. In addition, ERK3 promotes cancer cell growth and invasion in both kinase-dependent [3, 6] and kinase-independent mechanisms [5, 9]. On the contrary, it is largely unclear how ERK3 inhibits the growth and invasiveness of melanoma and hepatocarcinoma cells.

ERBBs are a family of structurally homologous receptor tyrosine kinases (RTKs), consisting of ERBB1 (also known as epidermal growth factor receptor (EGFR)), ERBB2, ERBB3, and ERBB4 [15]. ERBB RTKs are activated by a subset of growth factor ligands, including EGF and neuregulins (NRGs). Ligand binding induces receptor homo-dimerization or hetero-dimerization, followed by conformation change and kinase activation. ERBB2 has no known ligand binding, and ERBB3 lacks kinase activity, which makes them a preferable pair for heterodimerization in response to the binding of ERBB3 ligands, mainly NRGs such as NRG1 [16]. Activated ERBBs promote the activation of multiple downstream signaling pathways, primarily Ras/RAF/ERK1/2 and PI3K/Akt. ERBB RTK signaling is frequently upregulated in human cancers and plays critical roles in promoting tumor development and progression [15, 17].

Phosphatase and tensin homolog deleted on chromosome 10 (PTEN) is a dual lipid and protein phosphatase [18–19]. PTEN is a potent inhibitor of PI3K/Akt signaling and acts as a tumor suppressor in multiple human cancers [20]. Like many other tumor suppressor genes, PTEN is frequently dysregulated in cancers by genetic mutations (loss of function) and other molecular mechanisms, including downregulation of gene transcription and posttranslational modifications, leading to downregulation or even loss of protein expression and function. The roles of PTEN in tumor progression and metastasis have been studied in mice with tissue-specific deletion of PTEN [21]. For example, conditional deletion of PTEN in lung respiratory epithelial cells of bigenic mice containing both floxed PTEN alleles and a Cre recombinase transgene driven by a Clara cell secretory protein gene promoter (CCSP-Cre) causes bronchiolar hyperplasia [22], implying that another molecular alteration is required for lung tumor development within the context of PTEN loss.

Although recent studies have revealed important roles for ERK3 in promoting lung cancer cell growth in cultured cells and tumor growth in xenograft mouse models, it is unknown whether or not ERK3 plays a role in spontaneous lung tumorigenesis. To determine the role of ERK3 overexpression in lung tumorigenesis, we created a conditional ERK3 transgenic mouse line in which ERK3 transgene expression is driven by the ubiquitous *CAGGS* promoter and is controlled by Cre recombinase due to a floxed transcription STOP cassette inserted between the promoter and ERK3 transgene. By crossing with CCSP-Cre mouse line, we have found that while conditional ERK3 overexpression alone did not cause a clear phenotype in lungs, ERK3 overexpression cooperates with PTEN deletion to induce the formation of lung adenocarcinomas. Mechanistically, ERK3 overexpression stimulates activating phosphorylations of ERBB3 and ERBB2 by upregulating SP1-mediated NRG1 gene transcription.

## Results

### Conditional ERK3 overexpression and PTEN deletion induces tumorigenesis in mouse lungs

Previous studies from our lab and others’ have shown that ERK3 expression is upregulated in both lung adenocarcinomas (LUADs) and lung squamous cell carcinomas (LUSCs) of non-small cell lung cancers (NSCLC) and that ERK3 promotes lung cancer cell growth and invasiveness [3, 6]. One common limitation of previous analyses on ERK3 expression in NSCLC is the limited number of normal lung tissues in comparison with that of tumor samples. Hence we performed an analysis of ERK3 mRNA expression in NSCLCs (either LUADs or LUSCs) utilizing the GEPIA2 web server that analyzes differential gene expression in tumors versus a large number of normal samples from both the TCGA and GTEx projects [23]. As shown in Figure 1A, this analysis confirmed that ERK3 mRNA expression was upregulated in both LUSC and LUAD. In addition, by analyzing the c-Bioportal/TCGA NSCLC datasets [24] we found that high ERK3 expression level indicates poor overall survival of patients with lung adenocarcinomas (LUADs) (Figure 1B). These results suggest that ERK3 overexpression may promote NSCLC growth and progression. To test this, first we generated a conditional human ERK3 transgenic mouse line (LSL-ERK3) in which the CAGGS-LSL-huERK3 transgene (Figure S1A) was inserted specifically into Rosa26 gene locus. To validate the functionality of the transgene, LSL-ERK3 mouse line was crossed with the *Cre-ER™* line in which Cre expression is induced by tamoxifen treatment [25]. The littermates were genotyped by PCR for the expression of Cre and LSL-ERK3 transgenes (Figure S1B). Mice were then treated with tamoxifen for 5 days. As shown in Figure S1C, tamoxifen induced ERK3 transgene expression in lungs and livers.

**Figure 1.**
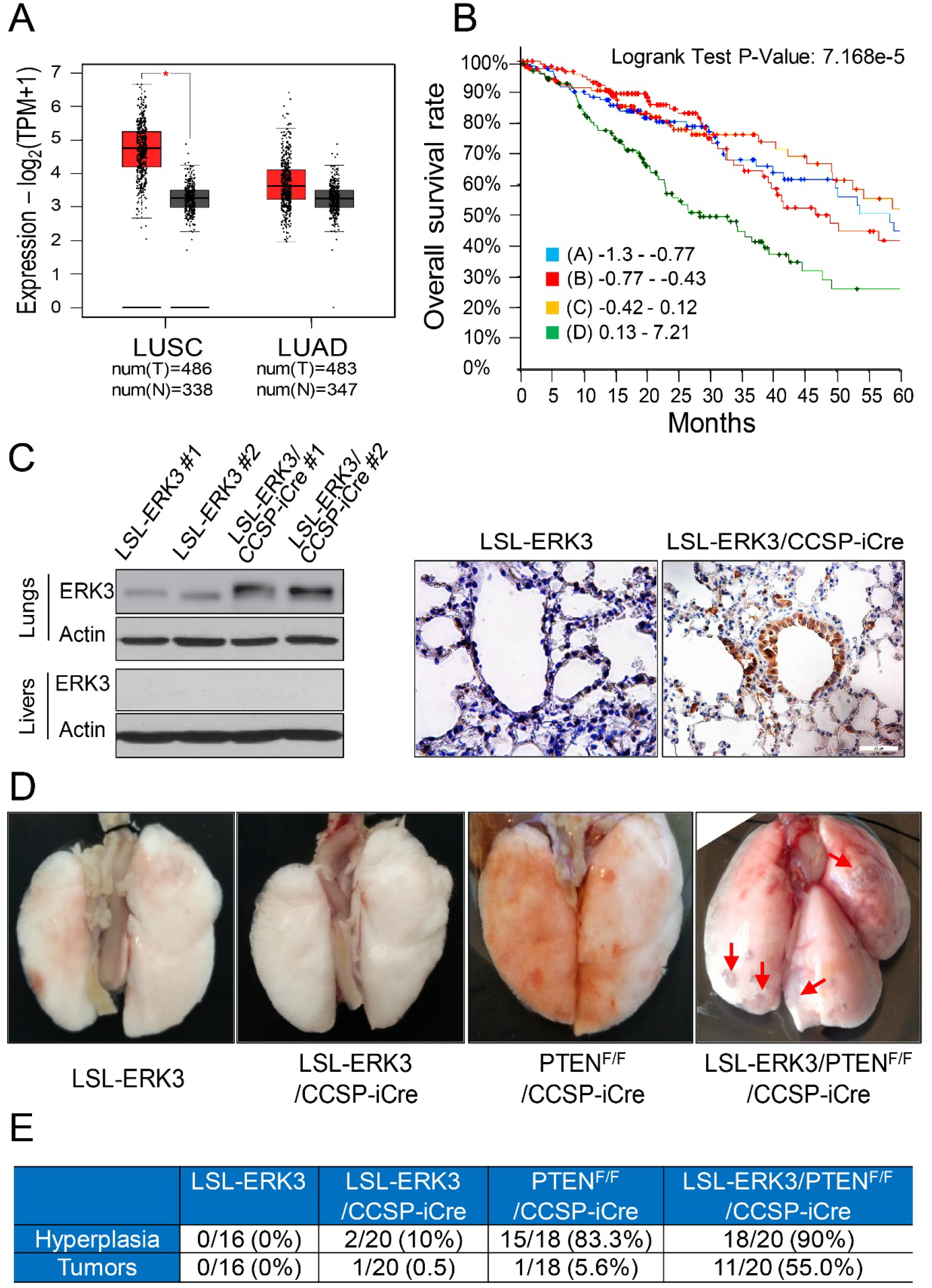
ERK3 overexpression induced lung tumor growth in PTEN-null background. (**A**) ERK3 (MAPK6) gene expression is upregulated in NSCLCs. Differential ERK3 mRNA expression in LUSCs or LUADs (from TCGA project) versus normal lung tissues (from both TCGA and GTEx projects) was performed using GEIPA2 web server. *: p <0.01, One-way ANOVA. num: indicates the total number of samples. T: indicates tumor. N: indicates normal tissues. (**B**) Lung adenocarcinoma patients (n=505, Data source: cBioportal/TCGA Lung adenocarcinoma Firehose legacy) were divided into 4 groups from quartiles of MAPK6 (ERK3) mRNA expression (Z scores relative to diploid samples, RNASeq V2 RSEM): group A (n=126; 42 decreased cases and median overall survival time: 58.41 months); group B (n=127; 41 decreased cases and median overall survival time: 46.68 months); group C (n=125; 36 decreased cases and median overall survival time: 66.59 months); group D (n=127; 64 decreased cases and median overall survival time: 28.38 months). (**C**) ERK3 overexpression is induced specifically in lungs by CCSP-iCre. ERK3 KI mouse was crossed with CCSP-iCre mouse to generate bigenic ERK3KI/CCSP-iCre mice. Cre-mediated overexpression of ERK3 protein in lungs of 1-year old bigenic ERK3KI/CCSP-iCre mice was first analyzed by Western blotting analysis (on the left) and then confirmed by immunohistochemical staining of ERK3 (images on the right). Livers were examined in Western blotting analysis to demonstrate the induction of ERK3 overexpression specifically in lungs. **(D**) Representative lungs of LSL-ERK3 mice, CCSP-iCre; LSL-ERK3 mice, CCSP-iCre; PTEN ^F/F^ mice and CCSP-iCre; LSL-ERK3; PTEN ^F/F^ mice at the age of one and a half years. As compared to LSL-ERK3 mice and CCSP-iCre; LSL-ERK3 mice, which have normal lungs, CCSP-iCre; PTEN ^F/F^ mice showed enlarged size of lung and CCSP-Cre; LSL-ERK3; PTEN ^F/F^ mice (D) have tumors on the surface of the lung (indicated by arrows). (**E**) Incidence of hyperplasia and formation of surface tumors in lungs of mice. The denominators and the numerators indicate the total number of mice analyzed in each group and the number of mice with tumor hyperplasia or tumor formation, respectively.

Having successfully generated the conditional ERK3 transgenic mouse line, we then crossed LSL-ERK3 mouse line with lung-specific CCSP-iCre line that expresses an improved Cre (iCre) inserted into the CCSP gene locus to determine whether overexpression of ERK3 specifically in the lung causes spontaneous lung tumor formation. While ERK3 protein overexpression was demonstrated by both Western blotting and immunohistochemistry specifically in the lungs of LSLERK3/CCSP-iCre mice (Figure 1C), no apparent phenotype was observed in the lungs of these mice (compare LSLERK3/CCSP-iCre with LSLERK3 control mice in Figure 1D and Figure 2A), indicating CCSP-iCre-induced ERK3 overexpression alone is insufficient for spontaneous lung tumorigenesis. Lung tumor formation usually requires multiple genetic alterations of both oncogenes and tumor suppressor genes [27]. *PTEN*, a tumor suppressor of the PI3K/Akt signaling pathway, is highly mutated in human lung cancer [28–29]. Since ERK3 overexpression alone did not induce lung tumors, we attempted to investigate the role of ERK3 overexpression in lung tumorigenesis under *Pten* deletion background. For this purpose, we generated triple transgenic mice that harbor LSL-*ERK3* transgene, *Pten* Floxed alleles (*Pten*^F/F^), and CCSP-iCre transgene (LSL-*ERK3*/*Pten* ^F/F^/CCSP-iCre, Figure S1D). As reported previously [22], CCSP-iCre-mediated *Pten* depletion induced lung hyperplasia but not tumor formation (CCSP-iCre/*Pten*^F/F^, Figure 1D). Importantly, tumors were observed on the surface of the lungs of LSL-*ERK3*/*Pten*^F/F^/CCSP-iCre mice (Figure 1D), and tumor incidence was about 50% (Figure 1E). These results suggest that conditional ERK3 overexpression cooperates with PTEN deletion to induce lung tumorigenesis.

**Figure 2.**
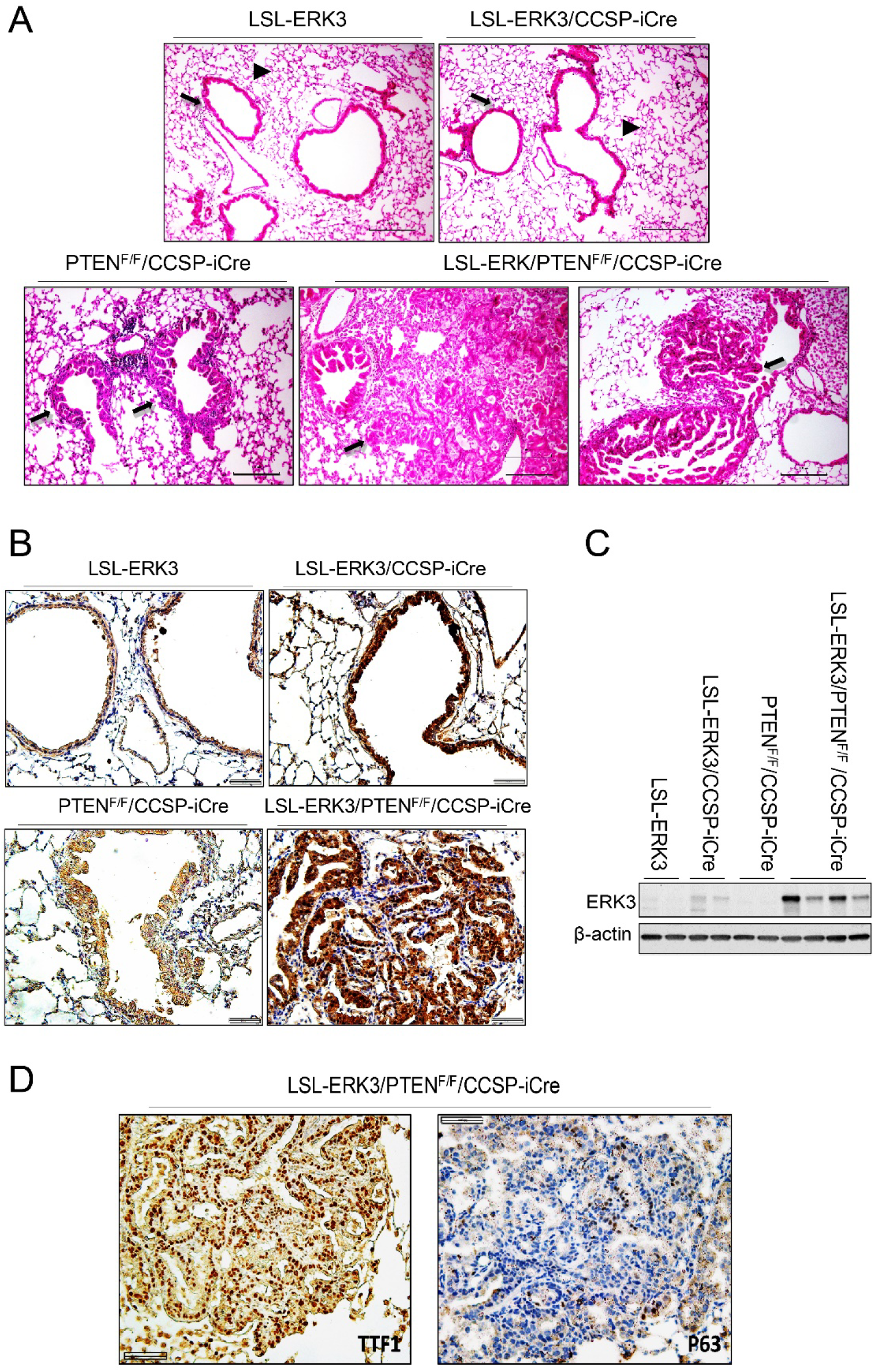
ERK3 overexpression induced lung adenocarcinoma development in PTEN-null background. (**A**) Hematoxylin and eosin (H/E) staining of formalin fixed and paraffin embedded (FFPE) lung sections of LSL-ERK3 and LSL-ERK3/CCSP-iCre mice, both of which display normal terminal bronchioles (indicated by arrows) and surrounding alveoli (indicated by arrow heads), PTEN^F/F^/CCSP-iCre displaying hyperplasia (indicated by arrows) of bronchiole epithelium, and LSL-ERK3/PTEN^F/F^/CCSP-iCre mouse displaying hyperplasia and tumors (indicated by arrows). (**B**) IHC of ERK3 protein expression in lungs of mice. (**C**) Western blotting analysis of ERK3 protein expression in lungs of mice. (**D**) IHC of TTF1 (a marker for LUAD) and P63 (a marker for LSCC) in the lungs of LSL-ERK3/PTEN^F/F^/CCSP-iCre.

### Concurrent ERK3 overexpression and PTEN deletion induce the formation of lung adenocarcinoma

There are two major subtypes of NSCLC: LUAD and LUSC on the basis of the pathological morphology and expression of biomarkers [26, 29]. To know which subtype(s) of lung tumors are formed in LSL-*ERK3*/*Pten*^F/F^/CCSP-iCre mice, we first performed histological analysis by H/E staining. As shown in Figure 2A, tumors appear to be acinar adenocarcinoma histologically. ERK3 overexpression in lung tumors was demonstrated by both IHC (Figure 2B) and Western blotting analysis (Figure 2C). We then confirmed the LUAD formation by IHC of biomarkers. Indeed, tumors in LSL-*ERK3*/*Pten*^F/F^/CCSP-iCre mice show prominent expression of TTF1 (a biomarker of LUAD) and faint staining of p63, a biomarker of LSCC (Figure 2C and Figure S2). These results suggest that concurrent ERK3 overexpression and PTEN deletion induce the formation of lung adenocarcinoma.

### ERK3 overexpression increases cell proliferation and reduces cell apoptosis in PTEN-null background

Next, we examined the effects of ERK3 overexpression on cell proliferation and survival in lungs. While conditional ERK3 overexpression alone did not show a clear effect on expression levels of Ki67 (a cell proliferation marker) in lungs (compare LSLERK3/CCSP-iCre with LSLERK3 control mice in Figure 3A and Figure 3B), ERK3 overexpression in the context of PTEN deletion greatly increased Ki67 expression level (compare LSL-*ERK3*/*Pten*^F/F^/CCSP-iCre with other groups in Figure 3A and 3B). These results suggest that ERK3 overexpression stimulates cell proliferation in PTEN deletion background. We then determined the effects on apoptosis by analyzing PARP cleavage and caspase 3 cleavage [30]. The levels of both cleaved PARP (Figure 3B right panel) and cleaved caspase 3 (Figure 3C) were greatly increased in the lungs of LSL-*ERK3*/*Pten*^F/F^/CCSP-iCre compared with other groups, suggesting concurrent ERK3 overexpression and PTEN deletion inhibits cell apoptosis. An increase in cell proliferation and a decrease in cell apoptosis account for the lung tumor formation in LSL-*ERK3*/*Pten*^F/F^/CCSP-iCre mice.

**Figure 3.**
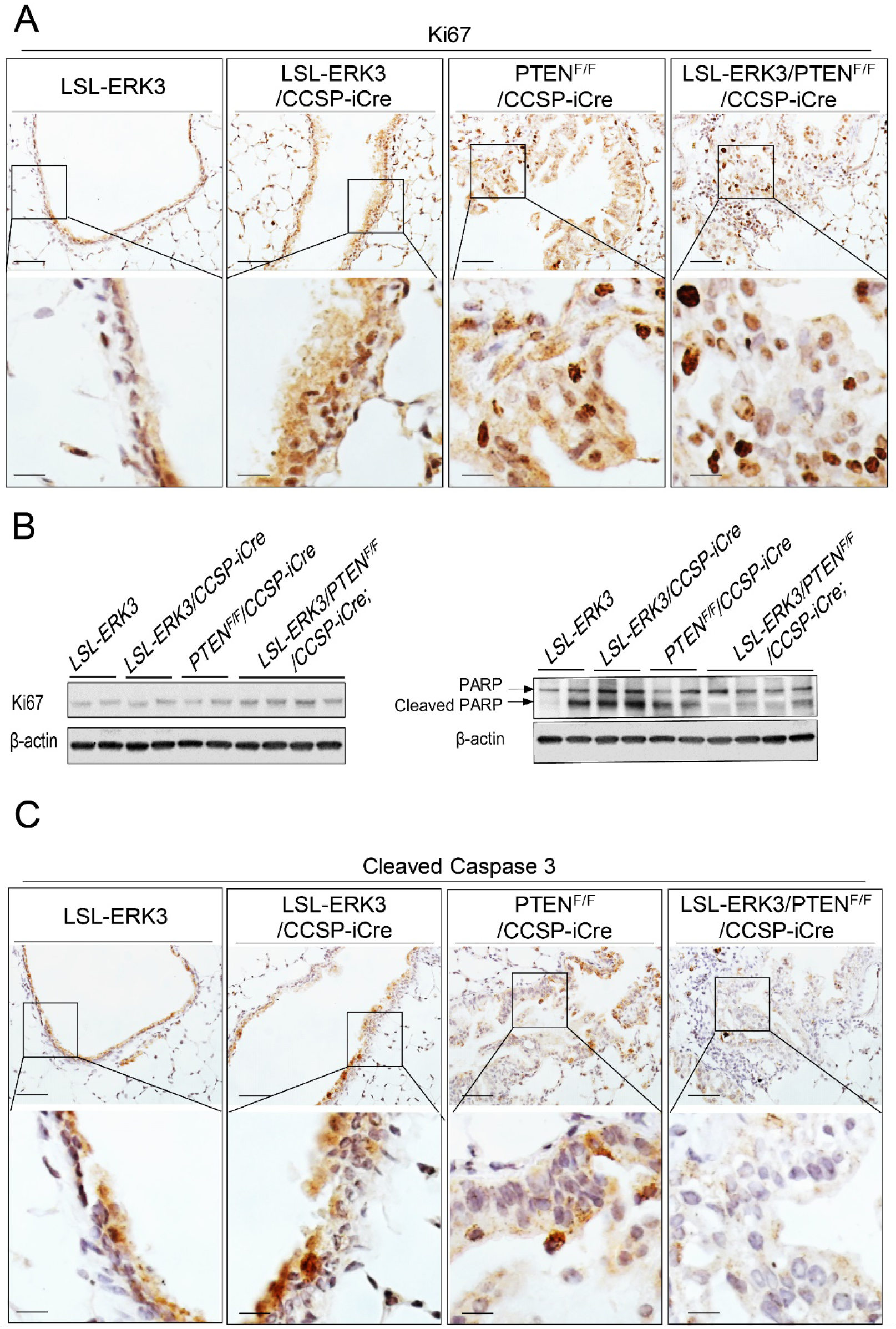
ERK3 overexpression increases cell proliferation and reduces cell apoptosis in PTEN-null background. **(A)** IHC of Ki67 protein expression in lungs of mice. (**B**) Western blot analyses of Ki67 and cleaved PARP levels in lungs of mice. (**C**) IHC of cleaved caspase 3 in lungs of mice.

### ERK3 overexpression increases activating phosphorylation of ERBB2 and ERBB3 in the context of PTEN deletion

ERBBs, in particular ERBB1, ERBB2, and ERBB3, play important roles in promoting lung tumor development and progression [31]. In an attempt to elucidate the molecular mechanism (s) by which ERK3 overexpression promotes tumor development, we examined the effects of ERK3 overexpression on the activating phosphorylation of ERBB1, ERBB2, and ERBB3. Interestingly, ERK3 overexpression in the context of PTEN deletion in LSL-*ERK3*/*Pten*^F/F^/CCSP-iCre mice greatly increased the levels of activating phosphorylation of ERBB3 and ERBB2, whereas it had little effect on ERBB1 (EGFR) phosphorylation (Figure 4A). As expected, Akt phosphorylation was greatly increased in PTEN^F/F^/CCSP-Cre mice (Figure 4B), but not further increased by ERK3 overexpression (compare LSL-*ERK3*/*Pten*^F/F^/CCSP-iCre with *Pten*^F/F^/CCSP-iCre mice, Figure 4B, and Figure S3). An increase in ERBB3 phosphorylation in LSL-*ERK3*/*Pten*^F/F^/CCSP-iCre was confirmed by immunostaining in the lungs (Figure 4C).

**Figure 4.**
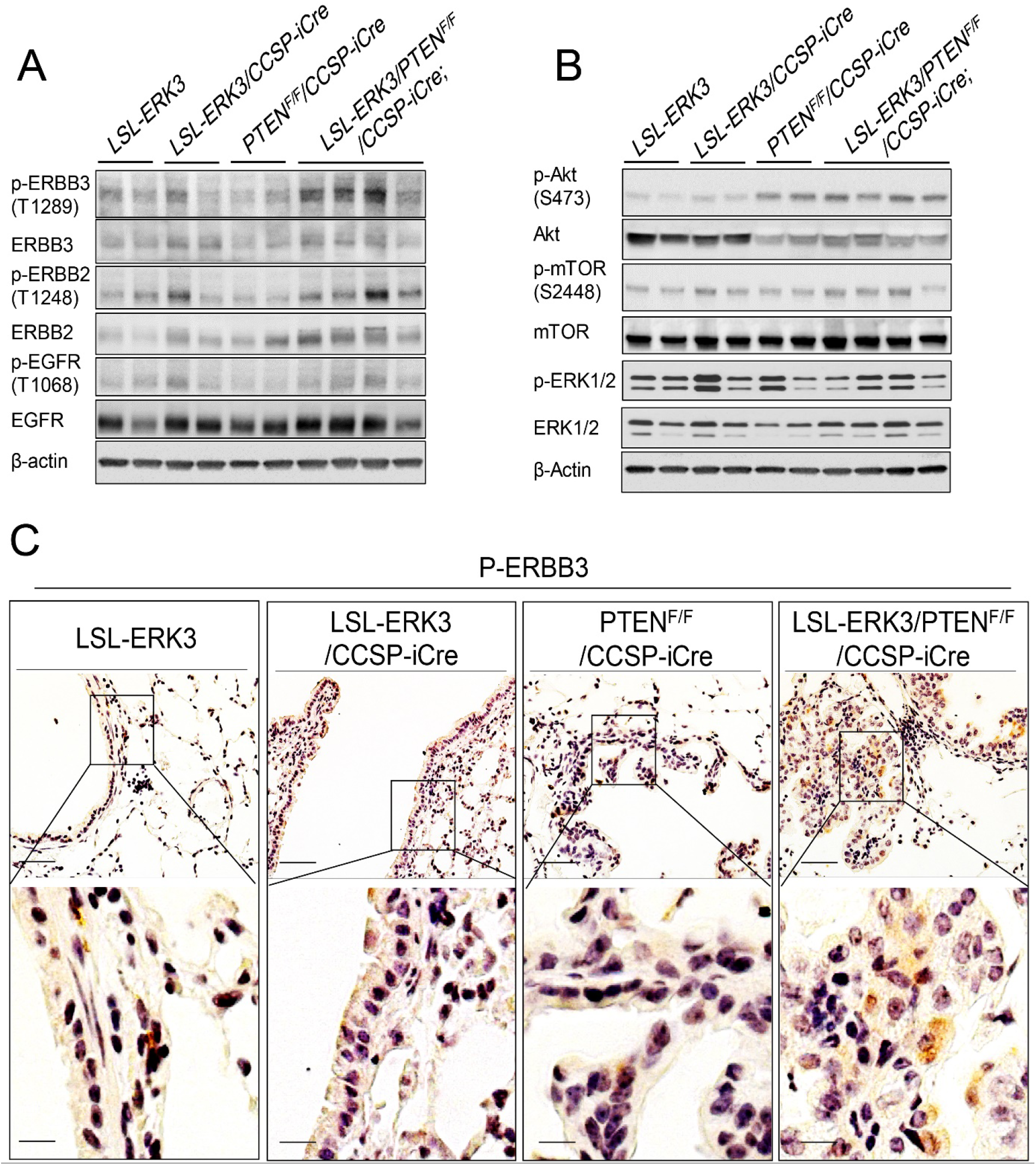
ERK3 overexpression increases the expression of P-ERBB3 in PTEN-null background. (**A**) Western blot analyses of activating phosphorylations of ERBBs (p-ERBBs) and total ERBB protein levels in lungs of mice. (**B**) Western blot analyses of activating phosphorylations and total protein levels of Akt, mTOR and ERK1/2 in lungs of mice. (**C**) IHC of p-ERBB3 in lungs of mice.

### ERK3 upregulates NRG1 gene transcript level in lung tumor cells

NRG1 is a major ligand for ERBB3 and induces heterodimerization and subsequent activation of ERBB3 and ERBB2 [16]. In addition, NRG1 was shown to be upregulated in NSCLC and stimulate NSCLC growth [32–33]. Thus, we determined whether ERK3 overexpression affected the NRG1 expression level, which may account for its effect in stimulating ERBB2/ERBB3 phosphorylation levels. Indeed, there is a significant increase in NRG1 transcript level in lung tumor tissues of LSL-*ERK3*/*Pten*^F/F^/CCSP-iCre mice as compared to *Pten*^F/F^/CCSP-iCre mice (Figure 5A), suggesting that ERK3 overexpression upregulates NRG1 gene expression in the context of PTEN deletion. The upregulation by ERK3 was confirmed in human lung cancer cell lines H520 and H1229, in which knockdown of ERK3 led to a significant decrease in NRG1 transcript levels (Figures 5B and 5C). SP1 transcription factor is known to bind to NRG1 gene promoter and regulate its transcription [34–35]. In addition, ERK3 was shown to stimulate SP1-mediated VEGFR2 gene transcription [36]. We therefore performed NRG1 gene promoter-driven luciferase assay for testing whether ERK3 coactivates SP1-mediated NRG1 gene transcription. Indeed, co-expression of SP1 and ERK3 synergistically stimulates NRG1 gene promoter activity in driving luciferase gene expression (Figure 5D).

**Figure 5.**
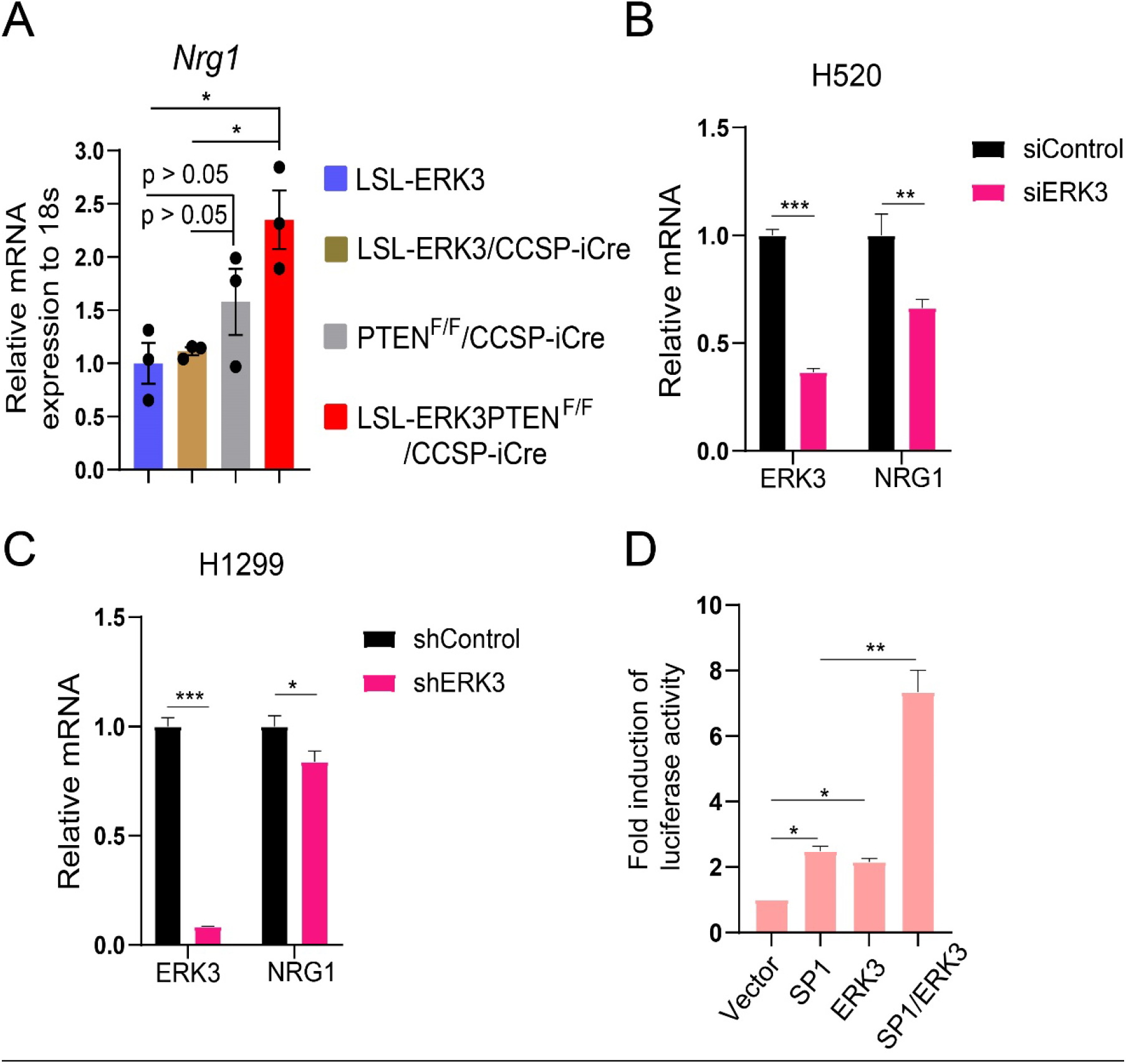
ERK3 upregulates NRG1 gene transcript level. **(A)** RT-qPCR analysis of NRG1 mRNA expression in lungs of mice. NRG1 mRNA expression level in mouse lungs of each different genotype was normalized to that of 18S and presented in relative to that of control LSL-ERK3 mouse lungs (arbitrarily set as 1). Results are expressed as mean ± SE of three independent experiments. * indicates P < 0.05 (Student’s *t*◻Jtest). (**B**) RT-qPCR analysis of NRG1 mRNA expression levels in H520 lung cancer cells treated with a siRNA specifically against ERK3 (siERK3) or a non-targeting control siRNA (siCtrl). Values represent mean ± SE of 3 independent experiments. Statistical significance was determined by Student’s t-test. **: p<0.01, ***: p<0.001. (**C**) RT-qPCR analysis of NRG1 mRNA expression levels in H1299 lung cancer cells stably expressing a shRNA specifically against ERK3 (shERK3) or a non-targeting control shRNA (shCtrl). Values represent mean ± SE of 3 independent experiments. Statistical significance was determined by Student’s t-test. *: p<0.05, ***: p<0.001. **(D**) ERK3 stimulates SP1-mediated NRG1 promoter activity. HeLa cells were co-transfected with pLightSwitch-NRG1 promoter-Luc and ERK3, SP1 or pSG5-empty vector control as indicated. The luciferase activity was measured 36 hours post◻Jtransfection using LightSwitch Luciferase Assay Kit (SwitchGear Genomics). Values in bar graphs present the fold induction of luciferase activity relative to the vector control. Results are expressed as mean ± SE of three independent experiments. * indicates P < 0.05 and **P < 0.01 (Student’s *t*◻Jtest).

## Discussion

In recent years, accumulating studies have suggested an important role for ERK3 in promoting tumor cell growth and invasion in a variety of cancers, including lung cancer. However, the role of ERK3 in spontaneous tumor growth in animal models has not been reported. In the present study, we have found that conditional overexpression of ERK3 in lungs cooperates with PTEN deletion to promote the formation of lung adenocarcinoma. To our knowledge, our study is the first revealing a bona fide tumor-promoting role for ERK3 in vivo using genetically engineered mouse models. Together with previous findings showing important roles of ERK3 in cultured cells and in xenograft lung tumor model [3, 6], our findings corroborate that ERK3 acts as an oncoprotein in promoting LUAD development and progression.

ERK3 mutations, including those in the kinase domain, have been reported in several types of cancer, but the frequency of these mutations is low [37, 38]. More frequent in cancers is the upregulation of ERK3 expression. Several studies, including those in TCGA, have shown the upregulation of ERK3 at both mRNA level and protein level in NSCLC, including both LUAD and LUSC [3, 6]. The kinase activity and cellular functions of ERK3 are positively regulated by phosphorylation of S189 in the activation motif, although it remains elusive regarding the upstream signal for stimulating S189 phosphorylation [5, 39–41]. Importantly, the level of S189 phosphorylation, which was determined by mass-spectrometry-based phosphoproteomic analyses in the study, was shown to be significantly elevated in lung adenocarcinomas [6]. These clinic findings suggest that altered ERK3 signaling in cancers is mainly caused by upregulation of expression level and posttranslational modifications rather than genetic mutations.

ERK3 plays differential roles in cell growth in different types of cancers. In NSCLCs, the role of ERK3 on cell growth appears to be affected by other molecular alterations in cells. For example, while ERK3 depletion had little effect on the growth of lung cancer cell lines H1299 and H1650 that express wild type *KRAS* [3, 6], it greatly reduced cell growth and/or anchorage-independent colony formation of *KRAS*^G12C^-positive H23 and H2122 NSCLC cell lines and xenograft tumor growth of Calu-1 cell line also expressing *KRAS*^G12C^ [6]. Although *KRAS*^G12C^-activated ERK1/2 signaling is important for the growth of H2122, Calu-1, and H23, ERK3 depletion does not affect MEK/ERK1/2 activity, and its role in promoting cancer cell growth is independent of the ERK1/2 pathway [6]. The molecular mechanisms underlying ERK3’s growth-promoting roles in these cells remain to be explored. Similarly, in our present *in vivo* transgenic mouse study, we found that ERK3 overexpression alone did not show an apparent effect on lung epithelial cell growth (cell proliferation and apoptosis data). However, in the context of deletion of PTEN tumor suppressor, ERK3 overexpression increased cell proliferation, decreased cell apoptosis, and promoted tumor formation. These findings suggest that ERK3 itself may not be able to transform normal epithelial cells, but is capable of promoting cancer cell growth and invasiveness once cells are transformed following the loss-of-function mutation of tumor suppressor gene or gain-of-function mutation(s) of oncogenes.

In contrast with the well-studied ERK1/2 signaling, little is known about the upstream stimuli and activators and downstream targets of ERK3. In an attempt to elucidate how ERK3 overexpression stimulates cell growth and tumorigenesis, we examined activating phosphorylation levels of ERBBs, ERK1/2, and Akt/mTOR, all of which are well-known oncogenic pathways in NSCLCs [31, 42]. Importantly, we found that conditional ERK3 overexpression in PTEN deletion background in lungs greatly increased the phosphorylation levels of ERBB3 and ERBB2 by upregulating their ligand NRG1 gene transcript level. Significant upregulation of NRG1 gene transcript was not seen in either ERK3 overexpression alone or PTEN deletion alone, suggesting that both ERK3 signaling and Akt signaling are required for stimulating NRG1 gene transcription in lung epithelium, which is likely mediated by SP1. NRG1 is a known target gene of SP1 transcription factor [34, 35]. Akt phosphorylates SP1, thereby stimulating SP1 transcriptional activity [43, 44]. In addition, SP1 transcriptional activity is also regulated by its coactivators such as SRC-3 [45, 46]. We have reported in our previous study that ERK3 phosphorylates SRC-3, which stimulates the interaction of SRC-3 with SP1 and their transcriptional activity in VEGFR2 gene transcription [36]. Similarly in the present study, we found ERK3 greatly increased SP1 transcriptional activity on NRG1 gene promoter. A major downstream target of NRG1/ERBB3/ERBB2 signaling is PI3K/Akt. However, we did not observe a clear concomitant increase of Akt phosphorylation with NRG1/ERBB3/ERBB2 activation in lungs of LSL-*ERK3*/*Pten*^f/f^/CCSP-iCre mice, likely in that Akt is constitutively activated/phosphorylated due to PTEN deletion.

Lung cancer can be originated from different cell types, such as type I and type II epithelial cells in the distal lung and Clara cells in the proximal airway [47]. Accordingly, cell-type-specific lung tumor models have been generated by utilizing either Clara cell-specific CCSP-Cre- or type II epithelial cell-specific SPC (surfactant protein C)-Cre-mediated expression of oncogenes (*e.g.*, *Kras*) or deletion of tumor suppressors (*e.g.*, *Trp53* and *Pten*) [48, 49]. Although ERK3 is expressed in both epithelial cells and Clara cells [50], the cell-autonomous functions of ERK3 in each of these cell types are unknown. Hence, the functions of ERK3 in lung tumor development and progression can be cell origin-dependent. Therefore, it would be important to further investigate the roles of ERK3 in lung tumor development and progression using other Cre-expressing systems such as using SPC-Cre mouse line or by intratracheal administration of Cre-expressing adenoviruses into the lungs for targeting multiple cell types [51].

## Materials and Methods

### Animal study

Animal work was done in accordance with protocols approved by the Animal Care and Use Committees of Baylor College of Medicine and Wright State University.

### Generation of conditional ERK3 transgenic mouse (LSLERK3)

Conditional ERK3 transgenic mouse was generated by using an established approach and following the procedures as previously described [52]. To generate an embryonic stem cell targeting construct, first, human ERK3 (huERK3) cDNA from pcDNA3-ERK3 plasmid [3] was subcloned by *Sal I* site into the shuttle vector RfNLIII (generously provided by Ming-Jer Tsai at Baylor College of Medicine, Houston, TX). Next, the fragment containing huERK3 cDNA and two homologous sequences for recombination with the base vector was released by KpnI/NheI digestion from the shuttle vector. The released DNA fragment and the targeting base vector were then electroporated into SW102 bacteria, subsequently leading to the generation of the targeting construct (pCAGGS-LSL-huERK3) through homologus recombination-based insertion of huERK3 into the targeting base vector upstream of the ubiquitous CAGGS promoter and downstream of a Lox-Stop-Lox (LSL) cassette. The targeting construct was verified by sequencing throughout the huERK3 cDNA and junction components.

The gene targeting in AB2.2 embryonic stem (ES) cells (mouse strain strain 129S5 background) and production of chimeras from those ES cells were performed by the Genetically Engineered Mouse Core at Baylor College of Medicine. Briefly, the targeting construct was linearized by PAC I digestion. The CAGGS-LSL-huERK3 transgene in the linearized targeting construct was specifically integrated through homologus recombination into Rosa 26 gene locus in AB2.2 ES cells. Chimeras were then produced using the correctly targeted ES clones. Chimeras were bred with C57BL/6 mice for the generation of the founder LSL-huERK3 transgenic mice. Germline transmission of the allele integrated with the LSL-huERK3 transgene was determined by mouse tail DNA PCR genotyping following the experimental conditions and procedures as described previously [52].

### Functional validation of conditional ERK3 transgene expression in LSL-ERK3 mouse

To validate the induction of ERK3 transgene expression by Cre protein, LSL-ERK3 mouse line was crossed with the *CAGG-Cre-ER™ mouse* line in which Cre expression is induced by tamoxifen treatment (JAX stock #004682) [25]. The littermates were genotyped by PCR for the expression of ERK3 transgenes and Cre following the procedures as described previously [53]. The littermates at the age of 5 weeks were administered with tamoxifen (75 mg/Kg body weight) once per day for a total of 5 consecutive days. The mice were sacrificed 3 days after the final injection, and lungs and livers were harvested for RNA and protein extraction.

### Generation of LSLERK3/CCSP-Cre, PTEN^F/F^/CCSP-Cre and LSLERK3/PTEN^F/F^/CCSP-Cre for lung tumorigenesis study

CCSP-iCre mouse [49] and floxed PTEN mouse (PTEN^F/F^) [54] were generated previously. LSLERK3 mouse was mated with CCSP-iCre to generate LSLERK3/CCSP-Cre mouse. LSLERK3/CCSP-Cre was then mated with PTEN^F/F^ to generate LSLERK3/PTEN^F/F^/CCSP-Cre mouse. Mice at different ages were sacrificed. Lungs were perfused using 1x PBS. The left lobe of lungs was then fixed by perfusion with 10% paraformaldehyde (PFA) for the use of histological analyses. The right lobes were frozen in liquid N2 and stored for later RNA or protein extraction.

### PCR genotyping

PCR genotyping using mouse tail DNA was performed following the experimental conditions and procedures as described previously (Wu et al). PCR primers are listed in table 1. The PCR conditions were: step 1: 15 seconds at 95°C; step 2: 40 seconds at 95°C for denaturation; step 3: 40 seconds at 56°C for annealing; step 4: 90 seconds of 72°C for elongation; step 5: repeating 33 cycles of steps 2-4; final step: hold at 4°C until use.

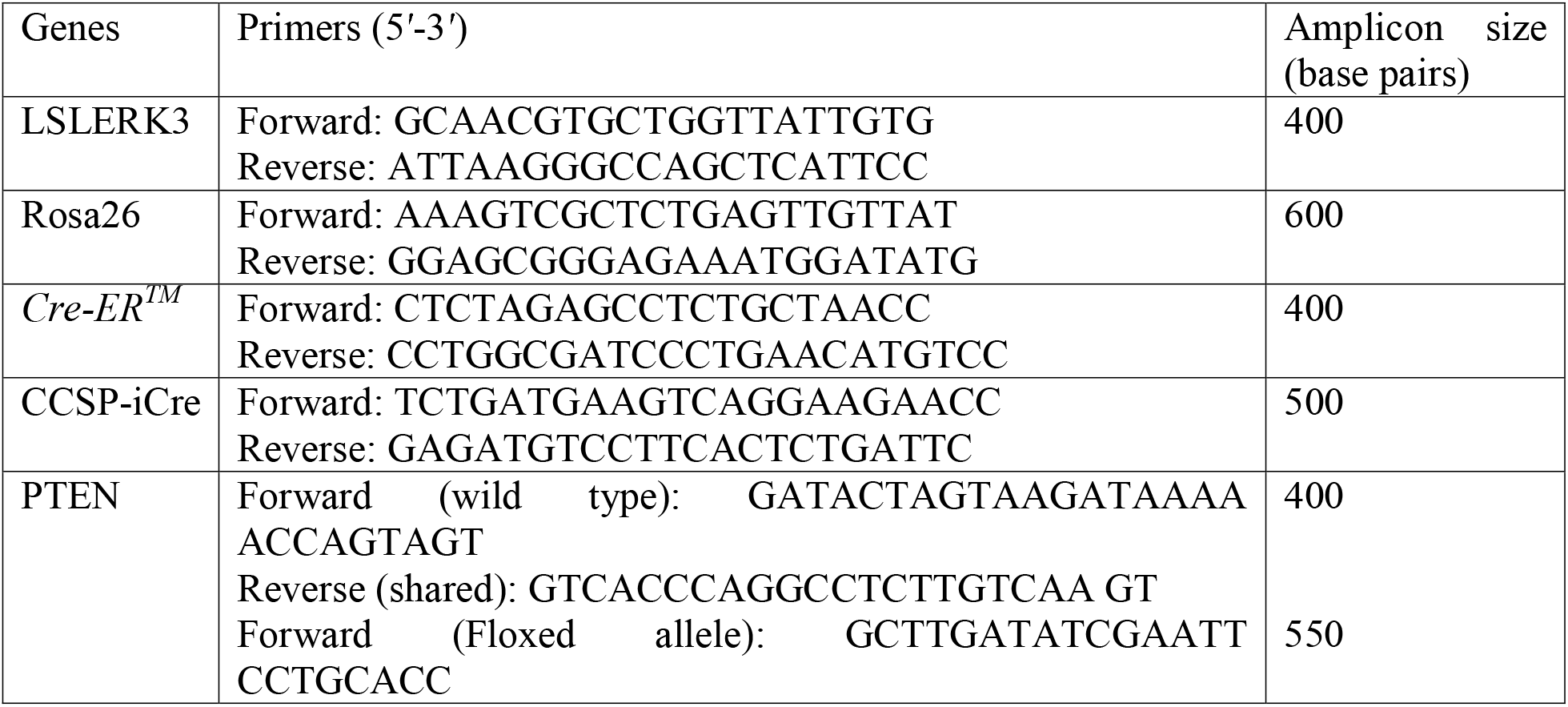

### Histopathology and immunohistochemistry

Histopathological analysis of PFA-fixed and paraffin-embedded lung tissues was performed by hematoxylin and eosin (H/E) staining and immunohistological staining following the procedures described in our previous study [49]. Briefly, for H/E staining, lung tissue sections (5 μm thickness) were dewaxed 3 times in xylene for 10 minutes each. Next, tissues were rehydrated for 5 minutes in each of the gradient ethanol concentrations (100%, 95%, 70%, 50%), followed by 5 minutes in double distilled water. The tissue sections were stained with hematoxylin [Vector Laboratories #H-3401] and eosin (Sigma Aldrich #SLBH6215V). For immunohistological staining, tissue slides were dewaxed and dehydrated as mentioned above. Antigen retrieval was then performed by treating the sections using Antigen unmasking agent (Vector Laboratories #H 3300) in an electric pressure cooker (Cuisinart-Model #CPC-600) for 15 minutes at high pressure setting. Thereafter endogenous peroxidase activity of the tissues was blocked by incubating the slides in 3% hydrogen peroxide in methanol for 10 minutes. Next, tissues were blocked in 5% normal goat serum (Vector Laboratories # s-1000) or MOM blocking reagents (Vector Laboratories, Cat# MKB-2213). Subsequently, sections were incubated with primary antibodies at 4°C overnight followed with biotinylated HRP-conjugated secondary antibody (Vector Laboratories # BA-1000) at room temperature for 1 hour. The slides were then developed using Vectastain ABC kit (Vector Laboratories # PK-6100) and diaminobenzidine (DAB, ACROS ORGANICS #112090250) substrate reagent (freshly prepared 1.7 mM DAB in 50 mM Tris (pH 7.6) containing 0.05% hydrogen peroxide) and then counter stained with hematoxylin. The primary antibodies used for immunohistochemistry are: anti-ERK3 (1:50 dilution, Abcam #ab53277), anti-TTF1 (1:1000 dilution, DAKO #M3575), anti-P63 (1:50 dilution, Santa cruz, # sc-8431), anti-Ki67 (1:5000, Abcam #ab15580), anti-cleaved caspase 3 (1:100; Cell Signaling Technology #CST9661), and anti-phospho-ERBB3 (1:100, Cell Signaling Technology # CST4791).

### Western blotting

Proteins were extracted from tissues, followed by Western blotting analysis following the procedures described previously [49]. Briefly, tissues were homogenized in EBC-lysis buffer containing 1 mM Complete protease inhibitors cocktail (Roche Diagnostics) and 1 mM protein phosphatase Inhibitor Cocktail I (Sigma Aldrich). Protein lysates were mixed with 5X SDS sample buffer and boiled then resolved on SDS-PAGE gels, followed by transfer onto nitrocellulose membrane and Western blotting. The Western blot was visualized by ECL chemiluminescence (Thermo Scientific). The primary antibodies used are: anti-ERK3 (Abcam #ab53277), anti-Ki67 (Abcam #ab15580), anti-PARP (Cell Signaling Technology, cat# 9532), anti-EGFR (Santa Cruz, SC-03), anti-Phospho-EGFR (Cell Signaling, CST3777), anti-ERBB2 (Santa Cruz, SC-284), anti-Phospho-ERBB2 (Santa Cruz, SC-293110), anti-ERBB3 (Santa Cruz, SC-285), anti-Phospho-ERBB3 (Cell Signaling, CST4791), anti-Akt (Cell Signbaling, CST4691), anti-phospho-Akt (Cell signaling, CST4060), anti-mTOR (Cell Signaling, CST2983), anti-phospho-mTOR (Cell signaling, CST2971), anti-ERK1/2 (Cell signaling, CST4695), anti-phospho-ERK1/2 (Cell Signaling, CST4370), and anti–β-actin (Sigma). β-actin was used as a loading control in Western blotting analysis.

### RNA extraction and Real Time Quantitative PCR (RT-qPCR)

Total RNA was extracted from cells using Trizol reagent (Thermo Scientific) and reverse transcription (RT) was done using SuperScript VILO Master Mix (Thermo Scientific) according to the manufacturer’s protocol. Quantitative PCR (qPCR) was performed using TaqMan Probe system (Roche Diagnostics) on the Applied Biosystems 7500 (Applied Biosystems) with either 18s RNA (for tissues) or GAPDH (for cells) as the internal control. Relative expression to normalizer sample was calculated using the ΔΔCT method.

### Cell culture and siRNA transient transfection

H1299 lung cancer cells stably expressing a shRNA specifically against ERK3 (shERK3) or a non-targeting control shRNA (shCtrl) were generated previously [3]. H520 lung cancer cell line and HeLa cervical cancer cell line were obtained from ATCC. H1299 and H520 were maintained in RPMI 1640 medium supplemented with 10% fetal bovine serum (FBS). HeLa cells were cultured in Dulbecco’s modified Eagle medium (DMEM) supplemented with 10% FBS. All the culture media and supplements were purchased from Life Technologies/Invitrogen. Transient transfection with siRNAs (20 nM working concentration) in H520 cells were done using DharmaFECT Transfection Reagent (Dharmacon) by following the manufacturer’s instructions. The silencer select siRNA targeting human ERK3 and the Silencer non-targeting Control #1 were purchased from Ambion.

### Luciferase reporter assay

pLightSwitch-NRG1 promoter-Luc (purchased from SwitchGear Genomics) is a luciferase expressing construct containing the human NRG1 gene promoter (927 bp fragment upstream of transcription start site). Plasmids pSG5-ERK3, pSG5-SP1 and the empty vector pSG5 were described in previous study [36]. HeLa cells were co-transfected with pLightSwitch-NRG1 promoter-Luc, pSG5-ERK3, pSG5-SP1 or pSG5-empty vector control using lipofectamine 3000 Reagent (Invitrogen). The luciferase activity was measured 36 hours post-transfection using LightSwitch Luciferase Assay Kit (SwitchGear Genomics).

### Statistics

Data are expressed as mean ± standard error (S.E.). Statistical significance was determined by a 2-sided Student’s *t*-test, where a p-value of less than 0.05 was considered statistically significant.

## Acknowledgments

This work was supported by a start-up fund of Wright State University and NCI 1R01CA193264-01 to Weiwen Long. We thank Dr. Ming-Jer Tsai, Dr. Sophia Tsai and Dr. San-Ping Wu at Baylor College of Medicine for the shuttle vector RfNLIII and the help with generating conditional ERK3 transgenic mouse line. We also thank Dr. Wei Wang at Baylor College of Medicine for breeding LSL-ERK3 mice and CCSP-iCre mice.

## Competing interests

The authors declare no competing interests.

## Author contributions

The experiments were designed by WL and JL. SV, JL, MM and WL carried out the experiments and data analysis. CCSP-iCre mouse line was generated by FD. The manuscript was written by WL and JL with inputs and comments from all coauthors.

---

**Supplemental Figure 1 (Figure S1).**
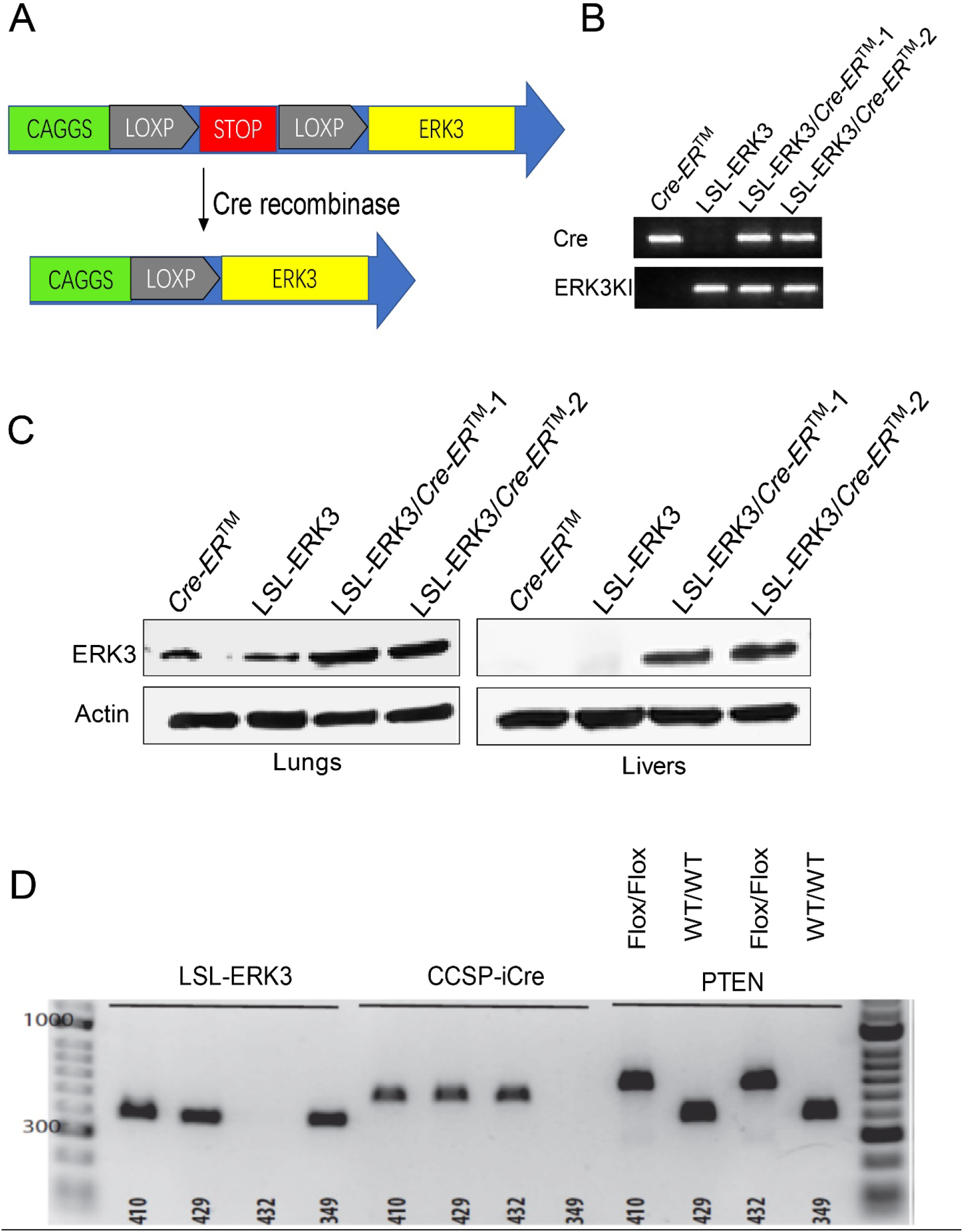
Generation of a transgenic mouse line conditionally expressing human ERK3. **(A)** Schematic illustration of conditionally controlled ERK3 transgene expression by Cre recombinase. Human ERK3 cDNA was cloned downstream of a ubiquitous CAGGS promoter, but was interspaced by a transcription STOP sequence that is flanked by two Lox P sites. The STOP sequence prevents the transcription of ERK3 driven by CAGGS promoter. In the presence of Cre recombinase, the STOP sequence flanked by Lox P sites will be excised and ERK3 expression will then be activated. The transgenic mouse line is designated as LSL-ERK3. **(B and C)** Characterization of LSL-ERK3 transgenic mouse line. LSL-ERK3 mouse was crossed with the CAGG-Cre-ER™ mouse line. The littermates were genotyped by PCR of tail DNA (B) for the presence of LSL-ERK3 and Cre transgenes. Mice at the age of 5 weeks were administered with tamoxifen (75 mg/Kg body weight) once per day for a total of 5 consecutive days. The mice were sacrificed 3 days after the final injection, and various tissues were harvested for RNA and protein extraction. Western blot analyses of ERK3 expression in the lungs and livers of different transgenic mice were shown in (**C**). IB: Immunoblot. (**D)** Representative PCR genotyping of experimental transgenic mice analyzed using tail DNA and specific primers for LSL-ERK3 (amplicon size: 400bp), CCSP-iCre (amplicon size: 450 bp), PTEN wild type allele (amplicon size: 400bp) and Floxed PTEN allele (amplicon size-450 bp). mouse #349: LSL-ERK3; mouse#432: PTEN^F/F^/CCSP-iCre; Mouse #429: LSL-ERK3/CCSP-iCre; mouse #410: LSL-ERK3/PTEN^F/F^/CCSP-iCre.

**Supplemental Figure 2 (Figure S2).**
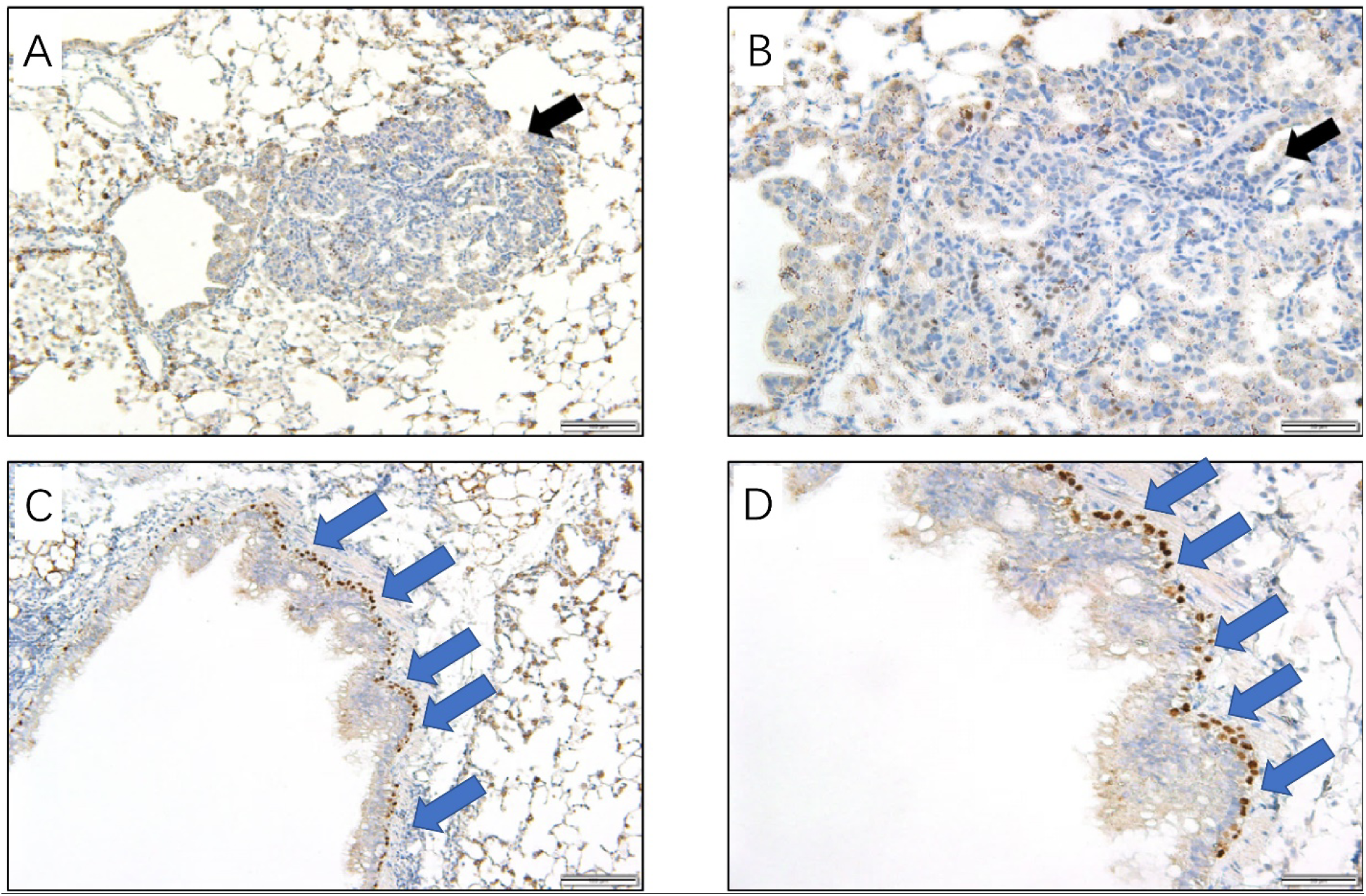
IHC staining of P63 in lung tumors and normal lung epithelium of LSL-ERK3/PTEN^F/F^/CCSP-iCre mouse. Prominent P63 staining is observed in normal epithelium (**C** and **D**) but not in tumors (**A** and **B**) of the same lung tissue section of LSL-ERK3/PTEN^F/F^/CCSP-Cre mouse. **A** and **C:** 20 X magnification. **B** and **D**: 40 X magnification.

## Notes

### Competing Interest Statement

The authors have declared no competing interest.

